## Supplementary Figures for "Characterizing transcriptomic responses to sediment stress across location and morphology in reef-building corals"

Supplementary materials for the manuscript “Characterizing transcriptomic responses to sediment stress across location and morphology in reef-building corals”

**Supplementary Figures**

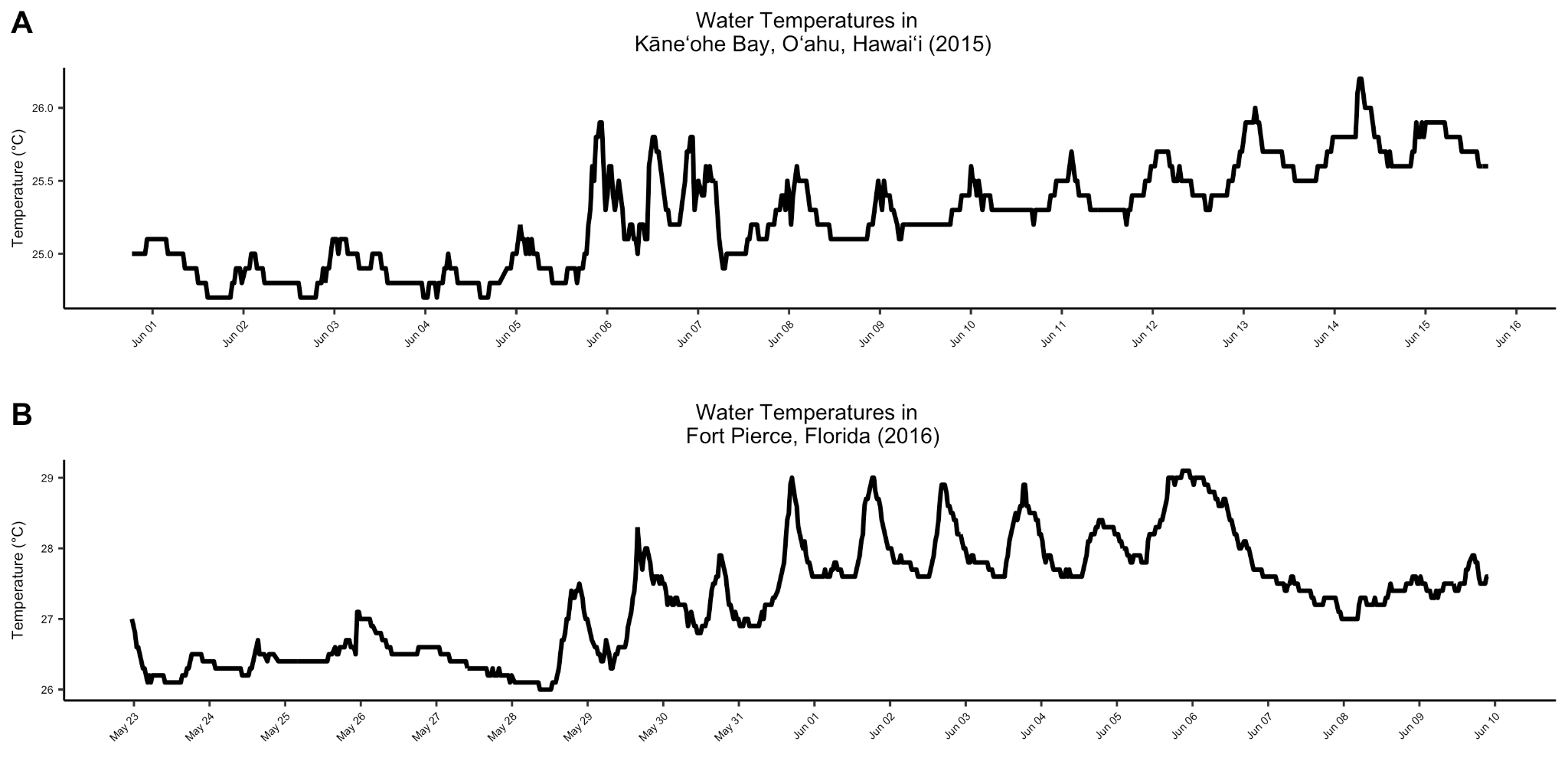

S1. (A) Sea surface temperature in Kāneʻohe Bay, Oʻahu, Hawaiʻi from June 1st to June 15th, 2015. Data obtained from Station 51207, Kāneʻohe Bay, HI (NOAA National Data Buoy Center). (B) Sea surface temperature in Fort Pierce, Florida from May 23rd to June 9th, 2016. Data obtained from Station 41114, Fort Pierce, FL (NOAA National Data Buoy Center).

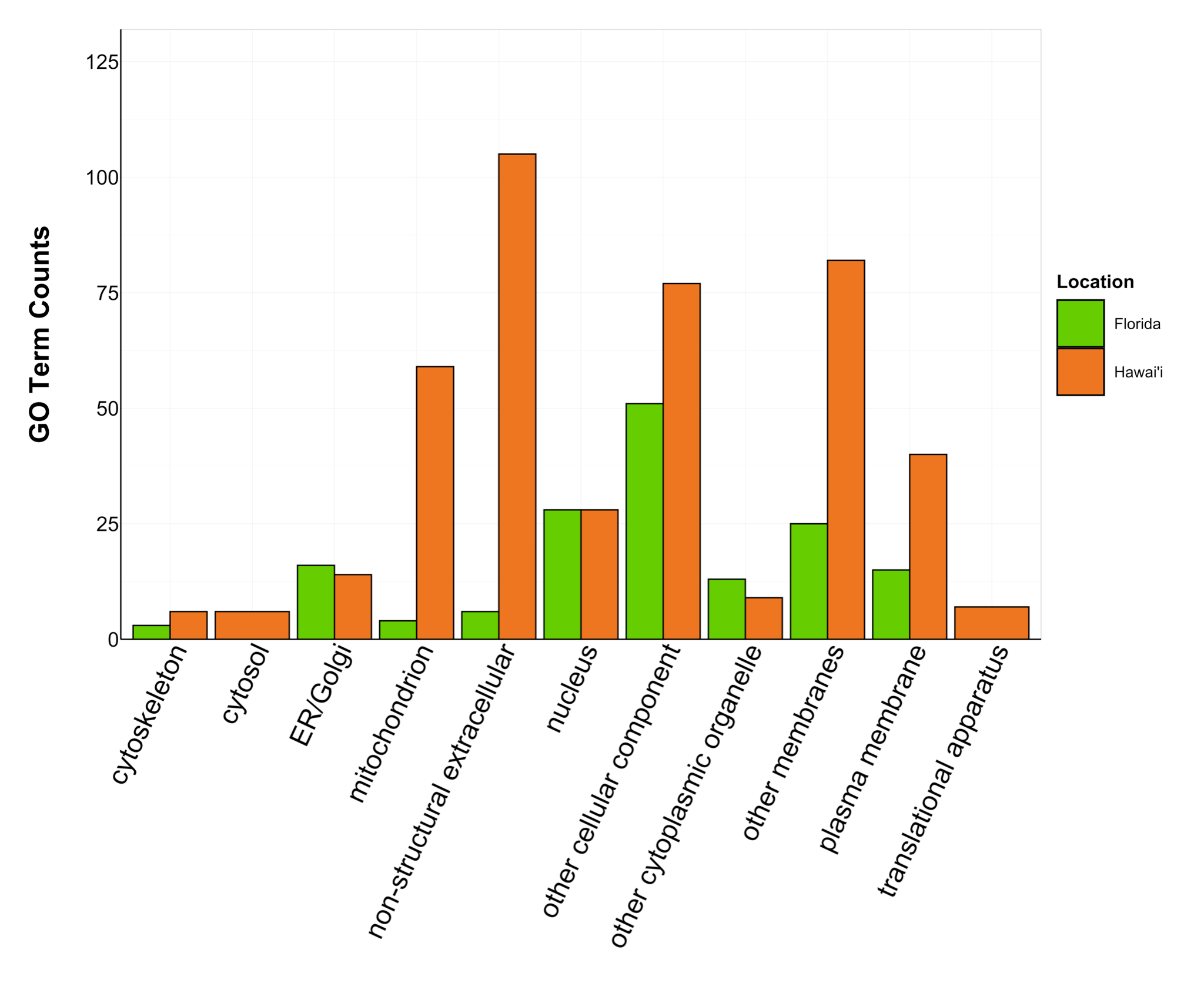

S2. Counts of Cellular Component GO terms grouped under GO slim categories by location. GO slim categories are on the x-axis, while the number of Cellular Component GO terms in each GO slim category is on the y-axis. The bars are colored by location: green bar = Florida, orange bar = Hawai’i.

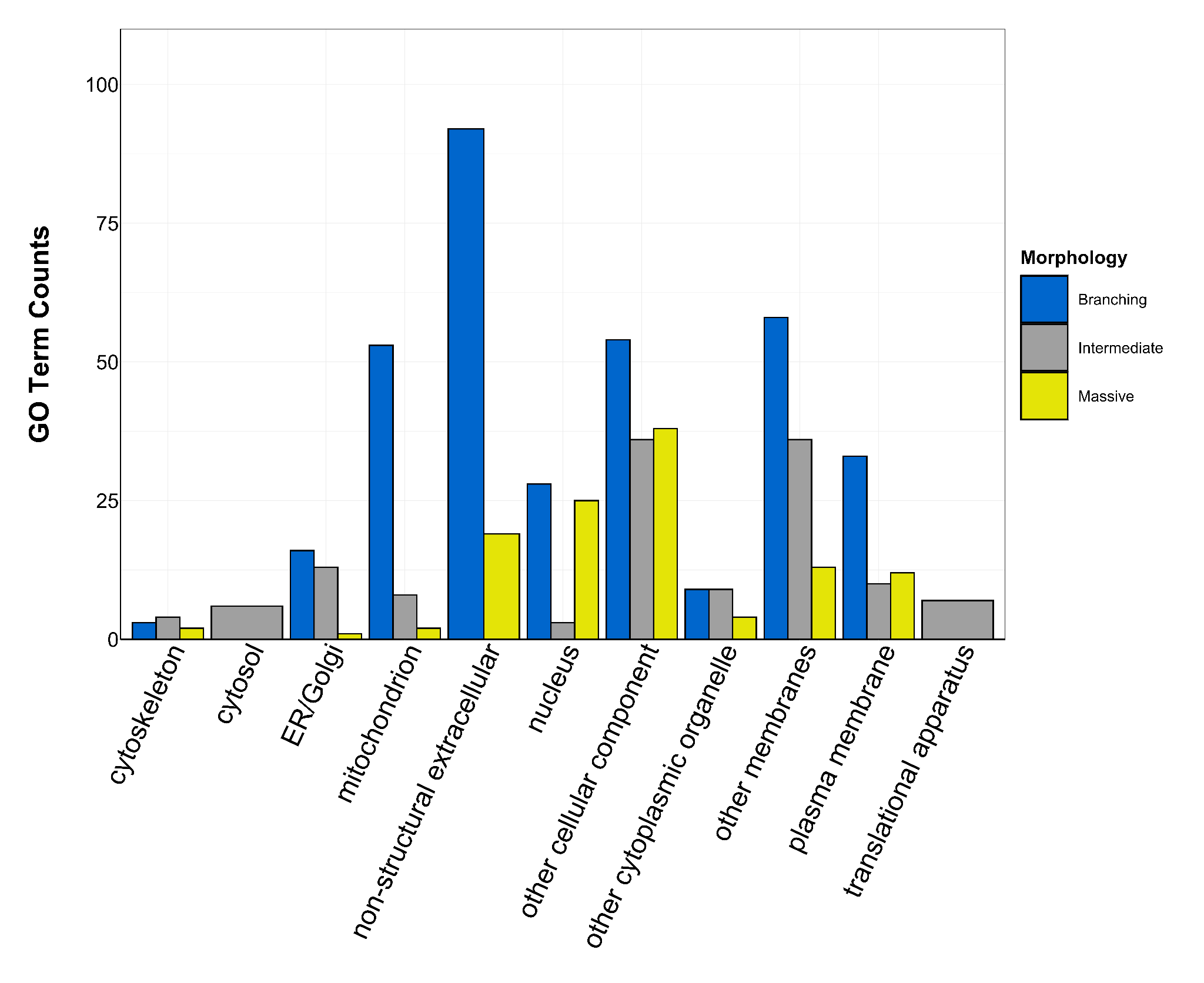

S3. Counts of Cellular Component GO terms grouped under GO slim categories by morphology. GO slim categories are on the x-axis, while the number of Cellular Component GO terms in each GO slim category is on the y-axis. The bars are colored by morphology: blue bar = branching, gray bar = intermediate, yellow bar = massive.

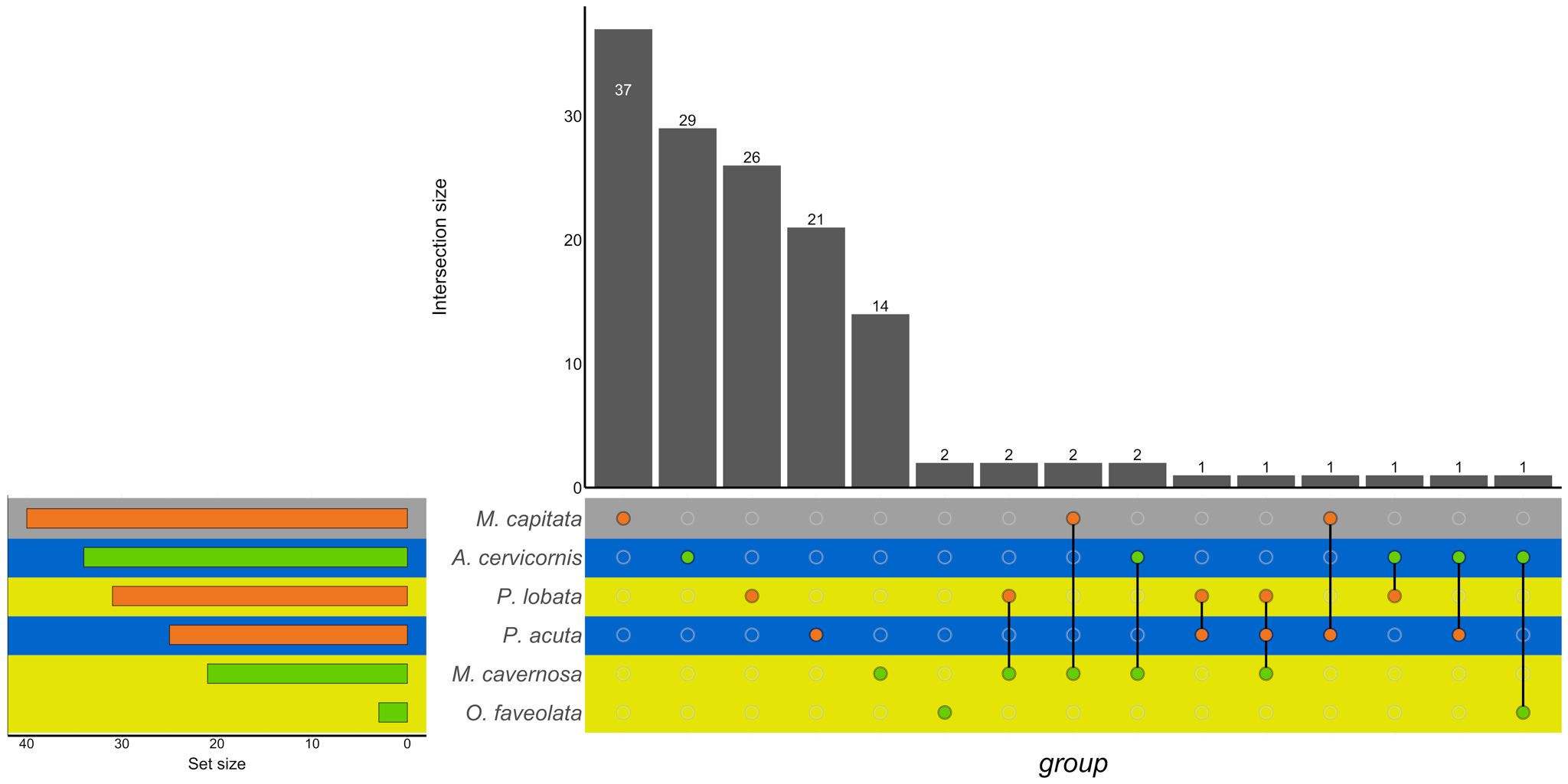

S4. UpSet plot of the intersection of Cellular Component GO terms across location (green bar or circle = Florida; orange bar or circle = Hawaiʻi) and morphology (blue stripe = branching; gray stripe = intermediate; yellow stripe = massive).

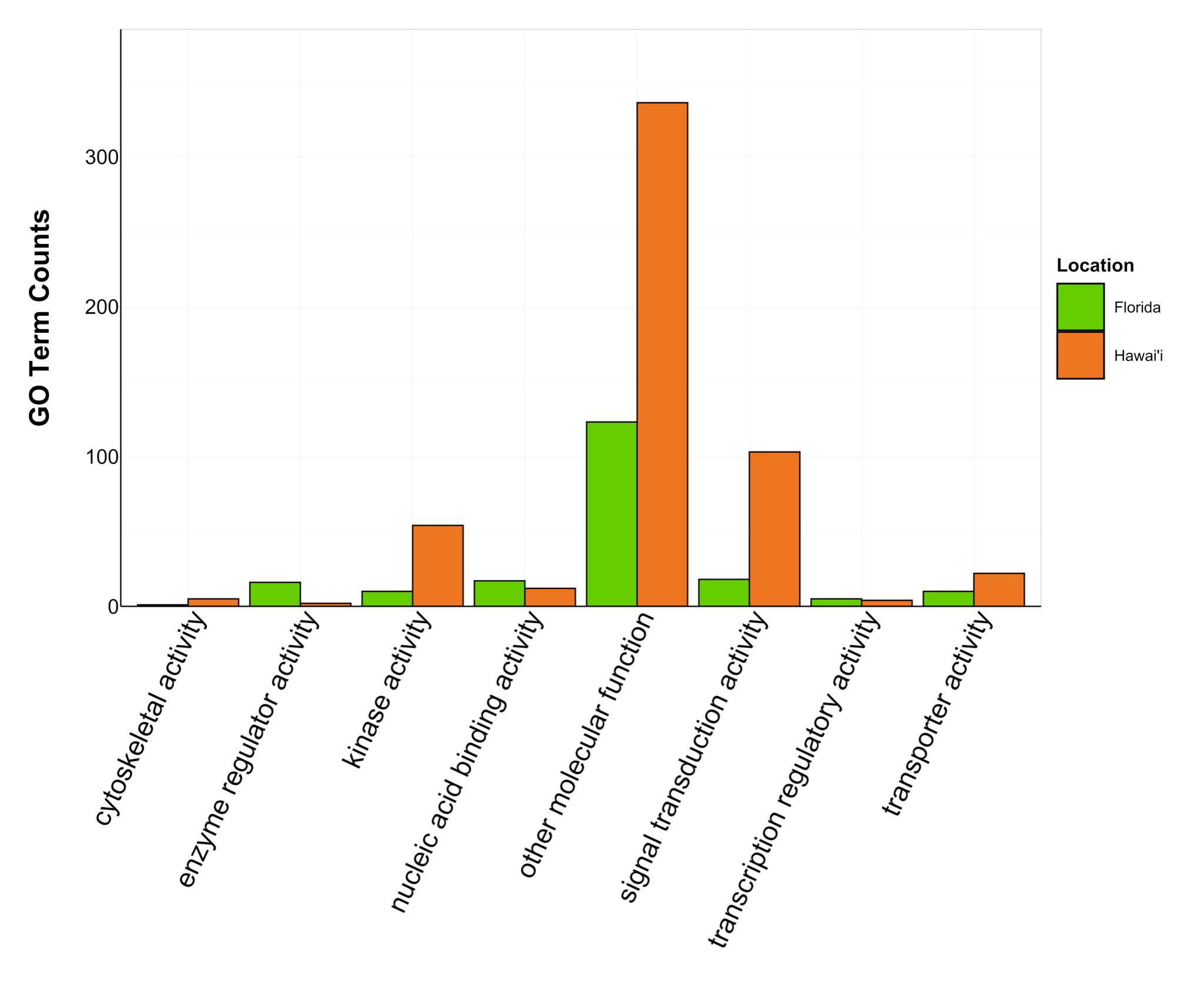

S5. Counts of Molecular Function GO terms grouped under GO slim categories by location. GO slim categories are on the x-axis, while the number of Molecular Function GO terms in each GO slim category is on the y-axis. The bars are colored by location: green bar = Florida, orange bar = Hawai’i.

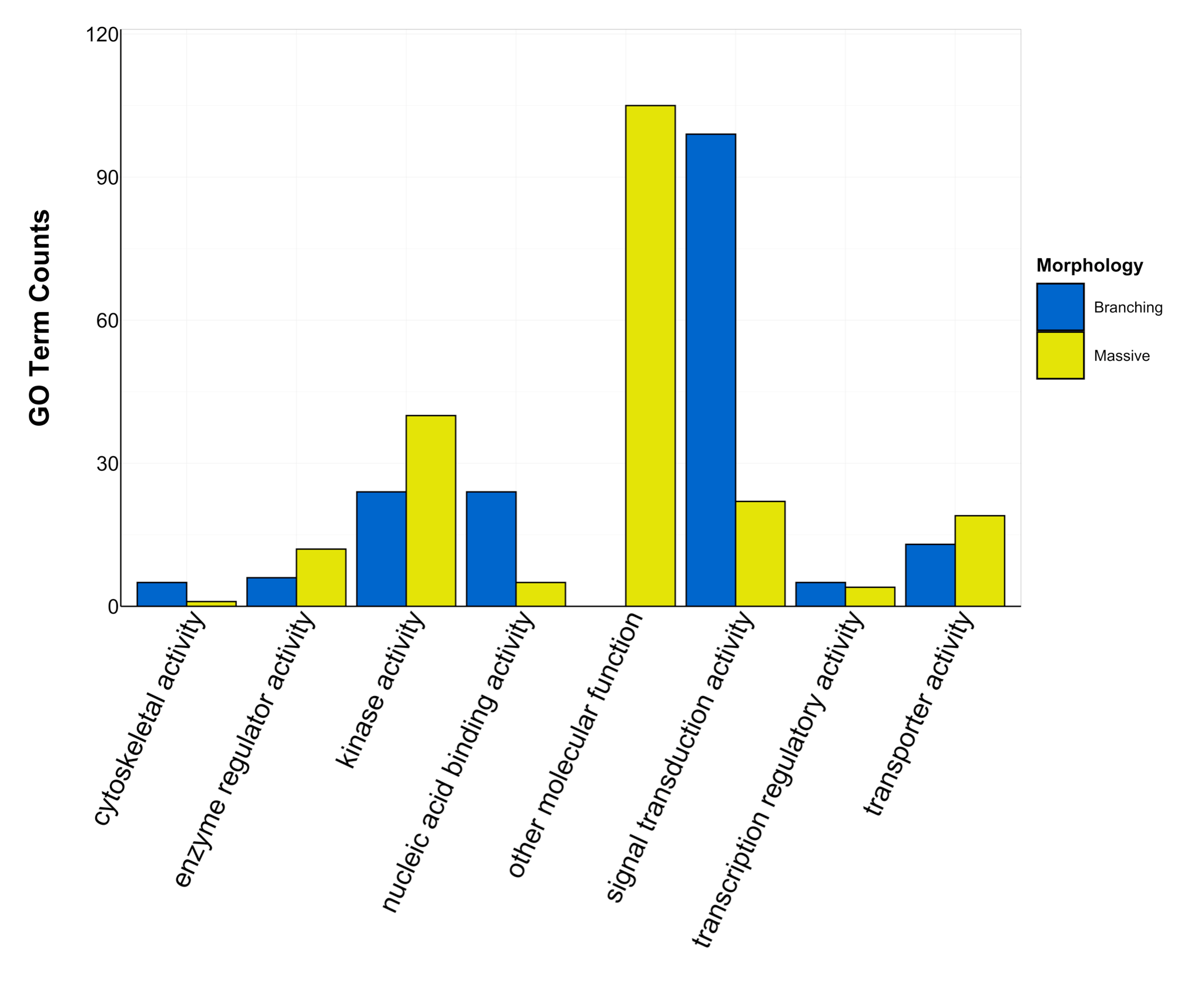

S6. Counts of Molecular Function GO terms grouped under GO slim categories by morphology. GO slim categories are on the x-axis, while the number of Molecular Function GO terms in each GO slim category is on the y-axis. The bars are colored by morphology: blue bar = branching, yellow bar = massive. Intermediate morphology was not assigned any Molecular Function GO terms.

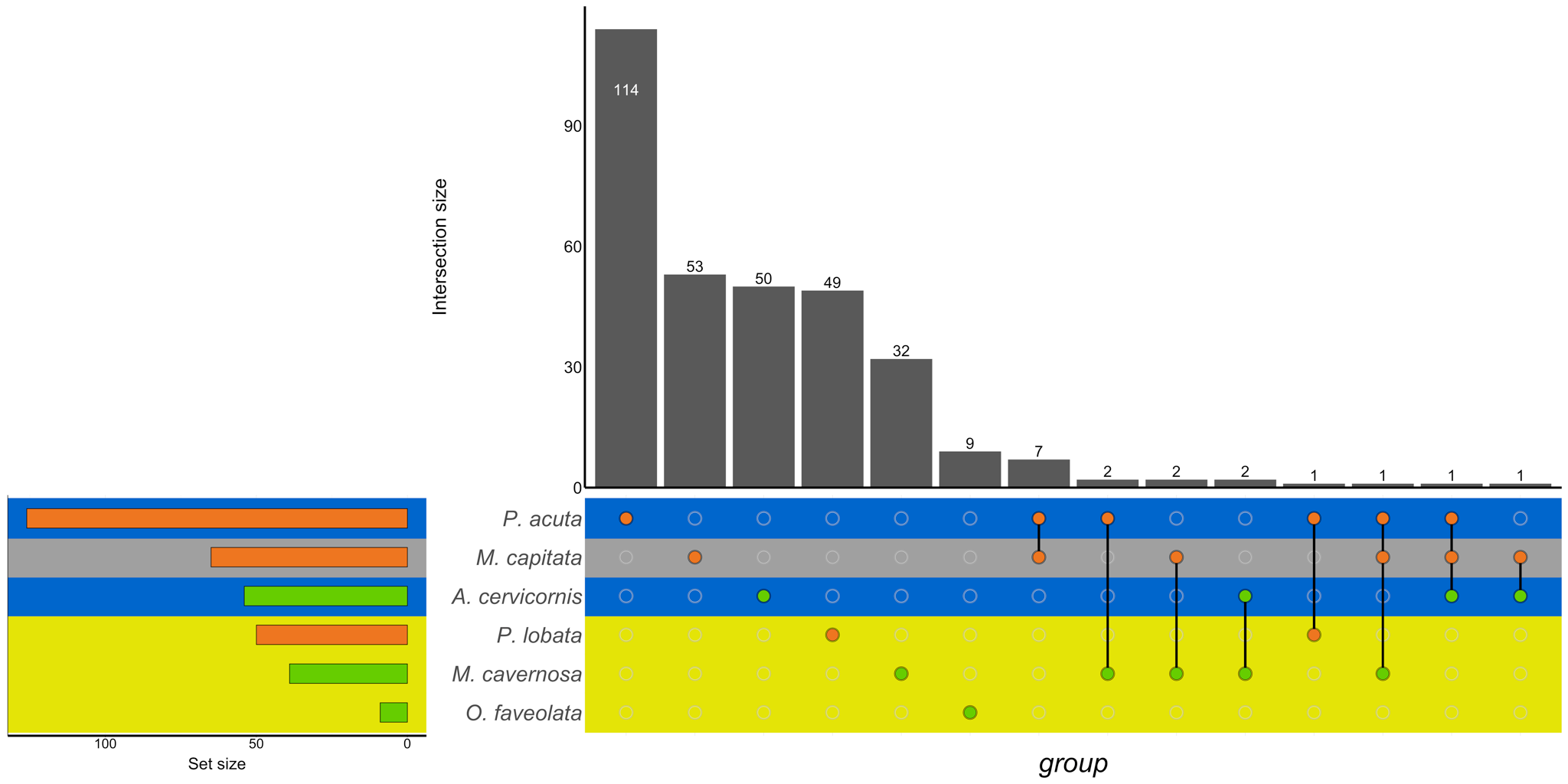

S7. UpSet plot of the intersection of Molecular Function GO terms across location (green bar or circle = Florida; orange bar or circle = Hawaiʻi) and morphology (blue stripe = branching; gray stripe = intermediate; yellow stripe = massive).
